# An Artificial Neural-Network Approach for Motor Hotspot Identification Based on Electroencephalography: A Proof-of-Concept Study

**DOI:** 10.1101/2021.05.09.443338

**Authors:** Ga-Young Choi, Chang-Hee Han, Hyung-Tak Lee, Nam-Jong Paik, Won-Seok Kim, Han-Jeong Hwang

## Abstract

**Background:** To apply transcranial electrical stimulation (tES) to the motor cortex, motor hotspots are generally identified using motor evoked potentials by transcranial magnetic stimulation (TMS). The objective of this study is to validate the feasibility of a novel electroencephalography (EEG)-based motor-hotspot-identification approach using a machine learning technique as a potential alternative to TMS.

**Methods:** EEG data were measured using 63 channels from thirty subjects as they performed a simple finger tapping task. Power spectral densities of the EEG data were extracted from six frequency bands (delta, theta, alpha, beta, gamma, and full) and were independently used to train and test an artificial neural network for motor hotspot identification. The 3D coordinate information of individual motor hotspots identified by TMS were quantitatively compared with those estimated by our EEG-based motor-hotspot-identification approach to assess its feasibility.

**Results:** The minimum mean error distance between the motor hotspot locations identified by TMS and our proposed motor-hotspot-identification approach was 0.22 ± 0.03 cm, demonstrating the proof-of-concept of our proposed EEG-based approach. A mean error distance of 1.32 ± 0.15 cm was measured when using only nine channels attached to the middle of the motor cortex, showing the possibility of practically using the proposed motor-hotspot-identification approach based on a relatively small number of EEG channels.

**Conclusion:** We demonstrated the feasibility of our novel EEG-based motor-hotspot-identification method. It is expected that our approach can be used as an alternative to TMS for motor hotspot identification. In particular, its usability would significantly increase when using a recently developed portable tES device integrated with an EEG device.

## Introduction

Motor impairment is a frequent symptom occurring after neurological disorders, such as stroke and Parkinson’s disease [1–3]. Although motor functions are not completely restored after neurological disorders, continuous rehabilitation is necessary to prevent muscle loss and the retrogression of intact motor functions. Conventional motor rehabilitation interventions involve specific movements related to the affected limbs, which are enforced by a therapist or an assistive rehabilitation device.

Recently, transcranial electrical stimulation (tES) capable of modulating cortical excitability using a weak electrical current has been introduced for motor rehabilitation, and its positive effects have been proven in many interventional studies even though the mechanisms have not yet been fully understood [4–10]. For example, one study showed that anodal transcranial direct current stimulation (tDCS) on the ipsilesional primary motor cortex could improve overall motor functions of the upper limbs in stroke patients, and its effects persisted for at least 3 months post-intervention [11]. Another study also demonstrated the positive effects of transcranial alternating current stimulation (tACS) on motor performance improvements in Parkinson’s disease [12].

To maximize the positive effects of tES on motor rehabilitation, it is important to find an optimal tES target location. The lesional primary motor area (M1) and its contralateral motor area have been traditionally used as stimulation target locations, to which anodal and cathodal tES electrodes are applied, respectively [13–15]. A cathodal tES electrode is sometimes applied to the contralateral supraorbital area of the lesional hemisphere instead of the contralateral motor area, called the M1-supraorbital prominence [15–17]. Identifying M1 can be performed by either the international 10-20 coordinate system used for electroencephalography (EEG) measurements or motor evoked potentials (MEPs), induced by transcranial magnetic stimulation (TMS). The former method uses C3 on the left hemisphere or C4 on the right hemisphere as the M1 location, depending on the lesional side [13, 18]. However, as the international 10-20 coordinate system does not consider inter-subject variability in cortical anatomy, the M1 location (C3 or C4) found by the international 10-20 coordinate system might not be an appropriate tES target location for motor rehabilitation. In particular, patients with neurological disorders, who are the main recipients of tES treatment, have shown significant changes in cortical morphologies owing to neural reorganization and plasticity after the occurrence of neurological diseases, such as stroke and cerebral palsy [19], requiring better methods to find individualized tES target locations more precisely.

As an alternative to the international 10-20 coordinate system, many studies have used TMS-induced MEP to find an individualized tES target location, called motor hotspot, for motor rehabilitation. This has been done because applying tES to the motor hotspot found by TMS-induced MEP could provide focal and accurate neuromodulatory effects on the motor network [20–22]. TMS is a useful tool to find motor hotspots, but requires a relatively bulky device and a somewhat cumbersome procedure accompanying the empirical judgment of a technician. Moreover, another device that measures MEP is required to find an individual motor hotspot using TMS.

In this study, we propose a novel alternative to TMS for motor hotspot identification based on EEGs measured during a simple finger-tapping motor task. A machine learning technique based on an artificial neural network (ANN) was applied to the measured EEG to localize individualized motor hotspots. The motor hotspot positions estimated by our proposed EEG-based machine-learning approach were then compared to those found by the traditional TMS-induced MEP to verify the feasibility of our approach. Preliminary results have been shown in [23].

## Methods

### Subjects

Thirty healthy subjects (10 females and 20 males; 25 ± 1.39 years; all right-handed) participated in this study, and they had no history of psychiatric diseases that might affect research results. All subjects received the information about the details of the experimental procedure and signed an informed consent for participation in the study. Appropriate monetary compensation for their participation was provided after the experiment. This study was approved by the Institutional Review Board (IRB) of Kumoh National Institute of Technology (No. 6250) and was conducted in accordance with the principles of the declaration of Helsinki.

### Motor Hotspot Identification by TMS-Induced MEP

Before measuring the EEGs, the motor hotspots of both hands were identified for each subject using the MEPs of the first dorsal interosseous (FDI) muscle. To this end, Ag-AgCl disposable electrodes were attached to the FDI muscles of both hands to measure the MEPs (actiCHamp, Brain Products GmbH, Gilching, Germany). We applied single-pulse TMS to the contralateral motor cortex during the resting state (REMED., Daejeon, Korea), and determined the motor hotspot location that showed the maximum MEP with individual minimum stimulation intensity [24–26]. The motor hotspot locations were marked in a 3D coordinate system based on the vertex (Cz in the international 10-20 system) using a 3D digitizer (Polhemus Inc., Colchester, Vermont, USA), and they were used as the ground truth to compare with those estimated by our EEG-based motor-hotspot-identification approach.

### EEG Recording

After identifying the individual motor hotspots using TMS for both hands, task-related EEG data were measured using 63 EEG electrodes attached to the scalp based on the international 10-20 system (Figure 1). The ground and reference electrodes were attached to Fpz and FCz, respectively. The positions of the EEG electrodes were marked in the 3D coordinate system as for the motor hotspot locations identified by TMS-induced MEPs. The EEG data were sampled at 1,000 Hz using a multi-channel EEG acquisition system (actiCHamp, Brain Products GmbH, Gilching, Germany) while the subjects were performing a simple finger-tapping motor task. Each subject performed the task 30 times, which consisted of pressing a button using their index fingers whenever a red circle was presented in the center of a monitor (Figure 2); two EEG measurement sessions were independently conducted using the left and right index fingers, respectively. The subjects were given sufficient rest whenever they wanted during the experiment to avoid fatigue. Moreover, they were instructed to remain relaxed during the experiment without any movement to prevent unwanted physiological artifacts. Because we carried out preliminary experiments with the first two subjects to validate our experimental paradigm before the main experiment, the right hand was only employed for these two subjects. In addition, we excluded the EEG data of one subject for both hands, and those of the left hand for other three subjects due to high contamination of EEG data caused by physiological artifacts. Thus, 29 and 25 EEG datasets were used for the right and left hands, respectively, for data analysis.

**Figure 1.**
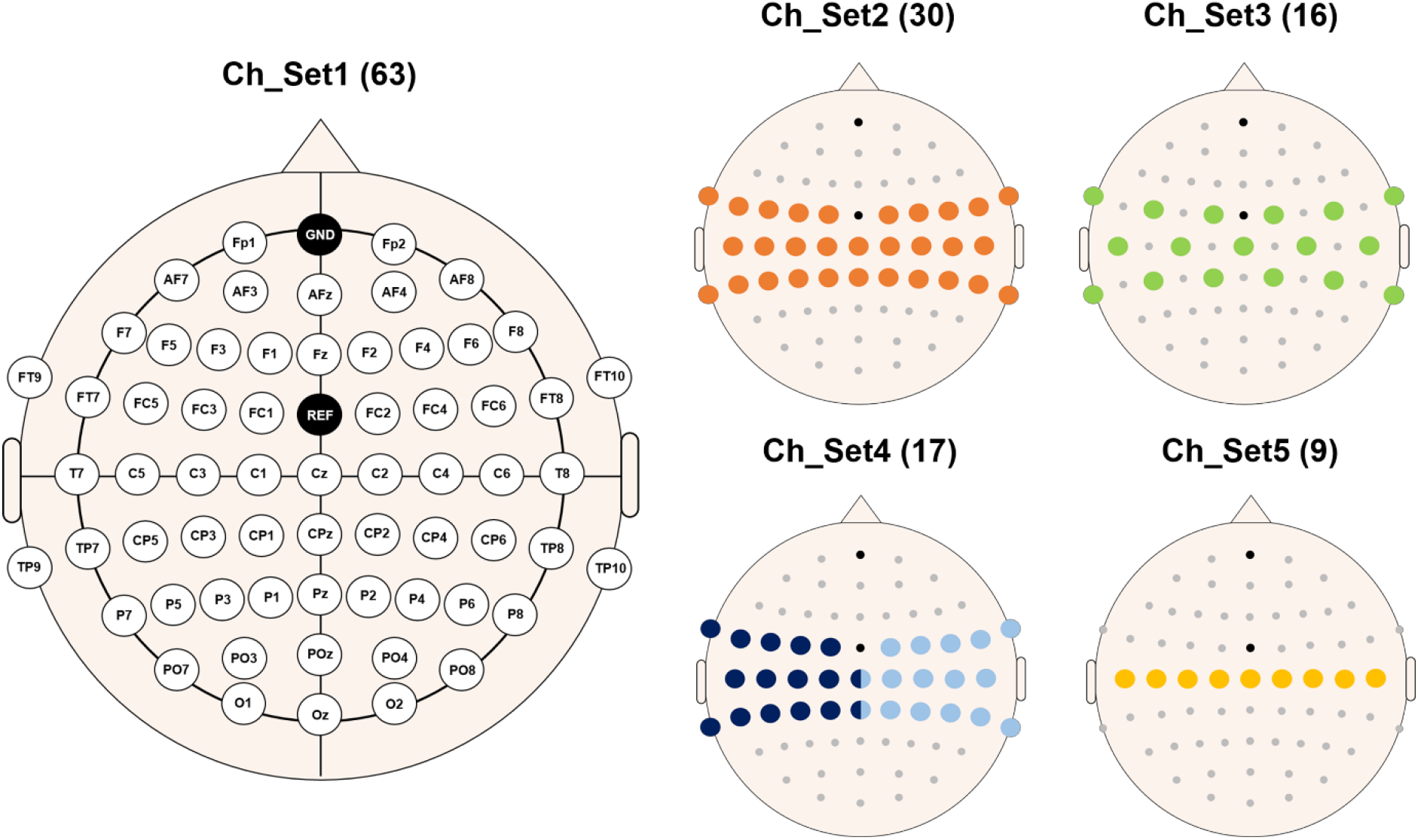
Electrode sites used for EEG data measurement. Five different channel sets were used for data analysis in order to study the impact of the number of channels on the error distance of motor hotspot location. Channels denoted by different colors in Ch_Set4 represent those selected on the contralateral motor cortex for the data analysis of each hand. The numbers in parentheses indicate the numbers of channels for the corresponding channel set.

**Figure 2.**
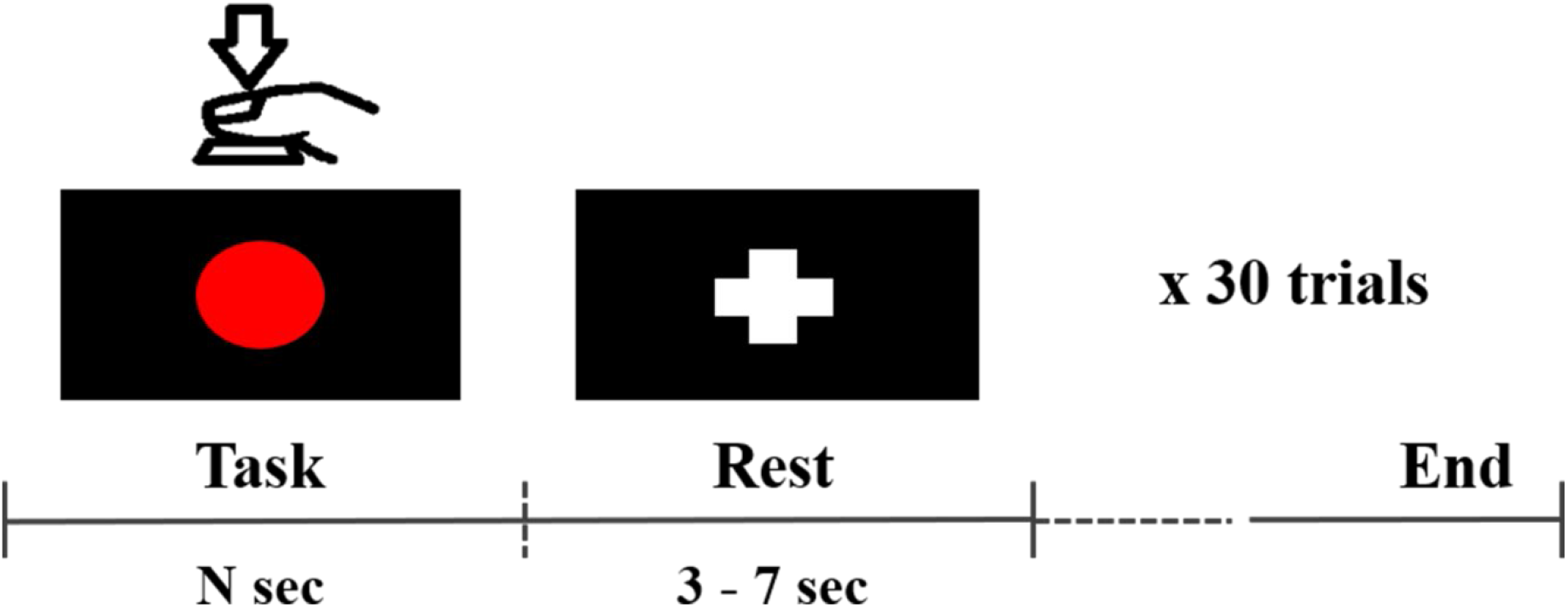
Experimental paradigm used in this study. Each subject presses a space bar whenever the red circle is presented in the middle of a screen. The red circle is maintained until the subject presses the space bar. After the task period, the fixation (‘+’) mark is presented to indicate a rest period.

### EEG-Based Motor Hotspot Identification

EEG signal preprocessing was performed using the EEGLAB toolbox based on MATLAB 2017b (MathWorks, Natick, MA, USA). The raw EEG data were first down-sampled into 200 Hz to which we sequentially applied common average reference and bandpass filtering between 0.5 and 50.5 Hz (zero-phase 3rd-order Butterworth filter). Independent component analysis (ICA) was then applied to the filtered EEG data to remove physiological artifacts. As mentioned above, from visual inspection we excluded the EEG data significantly contaminated, even after ICA application.

After preprocessing, we epoched EEG data between −0.5 and 0.5 s based on the key press point for each trial to extract motor-related EEG features. Power spectral densities (PSDs) of each EEG epoch were estimated in six frequency bands (delta: 1–4 Hz, theta: 4–8 Hz, alpha: 8–13 Hz, beta: 13–30 Hz, gamma: 30–50 Hz, and full: 1–50 Hz) using the fast Fourier transform (FFT). An ANN was used to estimate the motor hotspot location using PSD features where a subject-wise 10-fold cross-validation was performed with early stopping to prevent overfitting. The PSD features of each channel and the 3D location coordinates of the motor hotspot identified by TMS-induced MEP (ground truth) were used as the input for the ANN, and the 3D location coordinates estimated using the PSD features were used as the output. The numbers of input nodes, hidden nodes, and output nodes were 66 (63 channels + 3D motor hotspot coordinate), 40 (empirically selected), and 3 (3D coordinate estimated by ANN), respectively. We calculated the Euclidean distance between the 3D motor hotspot coordinates identified by TMS and EEG to quantitatively estimate the error distance of our EEG-based motor-hotspot-identification approach. The data analysis was independently performed for each of the six EEG frequency bands.

To investigate the impact of the number of channels on the error distance, we repeatedly performed the mentioned analysis by reducing the number of channels, in particular focusing on the channels attached to the motor area (Figure 1). For this investigation, only gamma-band PSD features (30–50 Hz) were used because they showed the best mean error distance when using all 63 channels.

## Results

Figure 3 shows the grand-average movement-related cortical potential (MRCP) measured during finger tapping between −0.5 to 0.5 s based on the task onset, which was obtained by averaging the MRCPs of four channels located on each hemisphere of the motor cortex to confirm the reliability of our EEG data (baseline period: −1 to −0.5 s). MRCP was clearly observed on the motor cortex during finger tapping for both hands, and, in particular, stronger MRCP was observed on the contralateral motor area [27–28].

**Figure 3.**
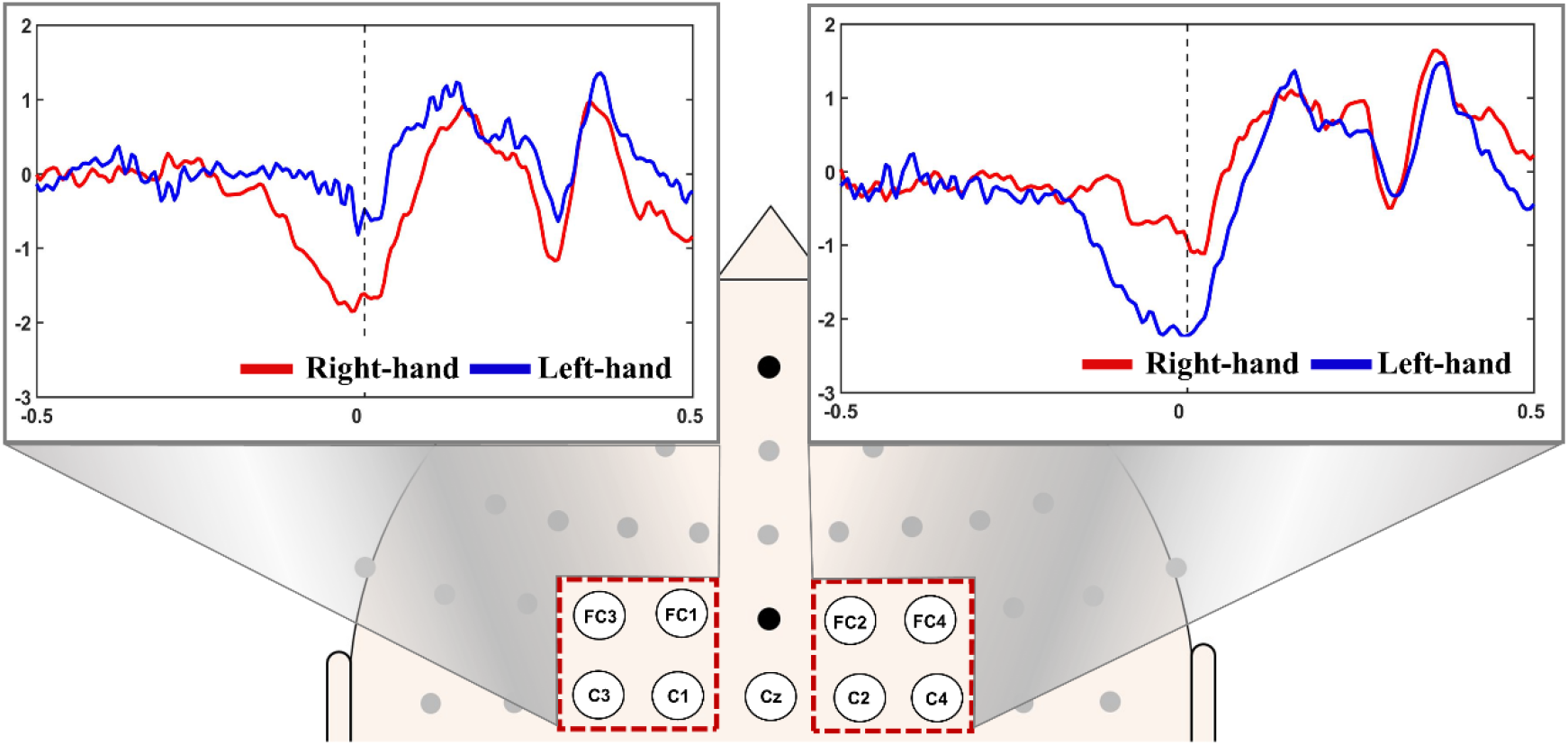
Grand averaged movement-related cortical potential (MRCP) patterns of all subjects on both hemispheres. Contralateral MRCPs are clearly observed.

Figure 4(a) shows the mean error distances of the motor hotspot locations estimated by our EEG-based machine learning approach for each hand with respect to the frequency band when using all 63 channels. The PSD features of the relatively high frequency bands (beta, gamma, and full bands) resulted in significantly lower mean error distances than those of the low frequency bands (delta, theta, and alpha) for both hands (RM-ANOVA with Bonferroni corrected *p*-value < 0.05: delta = theta = alpha > beta = gamma = full). No significant difference was observed between the left and right hand for all the frequency bands in terms of the error distance (paired t-test *p* > 0.05), except for the full band (right hand > left hand). Figure 4(b) shows a representative example of a single subject displaying the 3D locations of the motor hotspot identified by TMS (blue rectangle) and those of our EEG-based approach for both hands with respect to the frequency band. The motor hotspots estimated using the relatively high frequency bands were found closer to the ground truth motor hotspot identified by TMS, as compared to those estimated using the low frequency bands.

**Figure 4.**
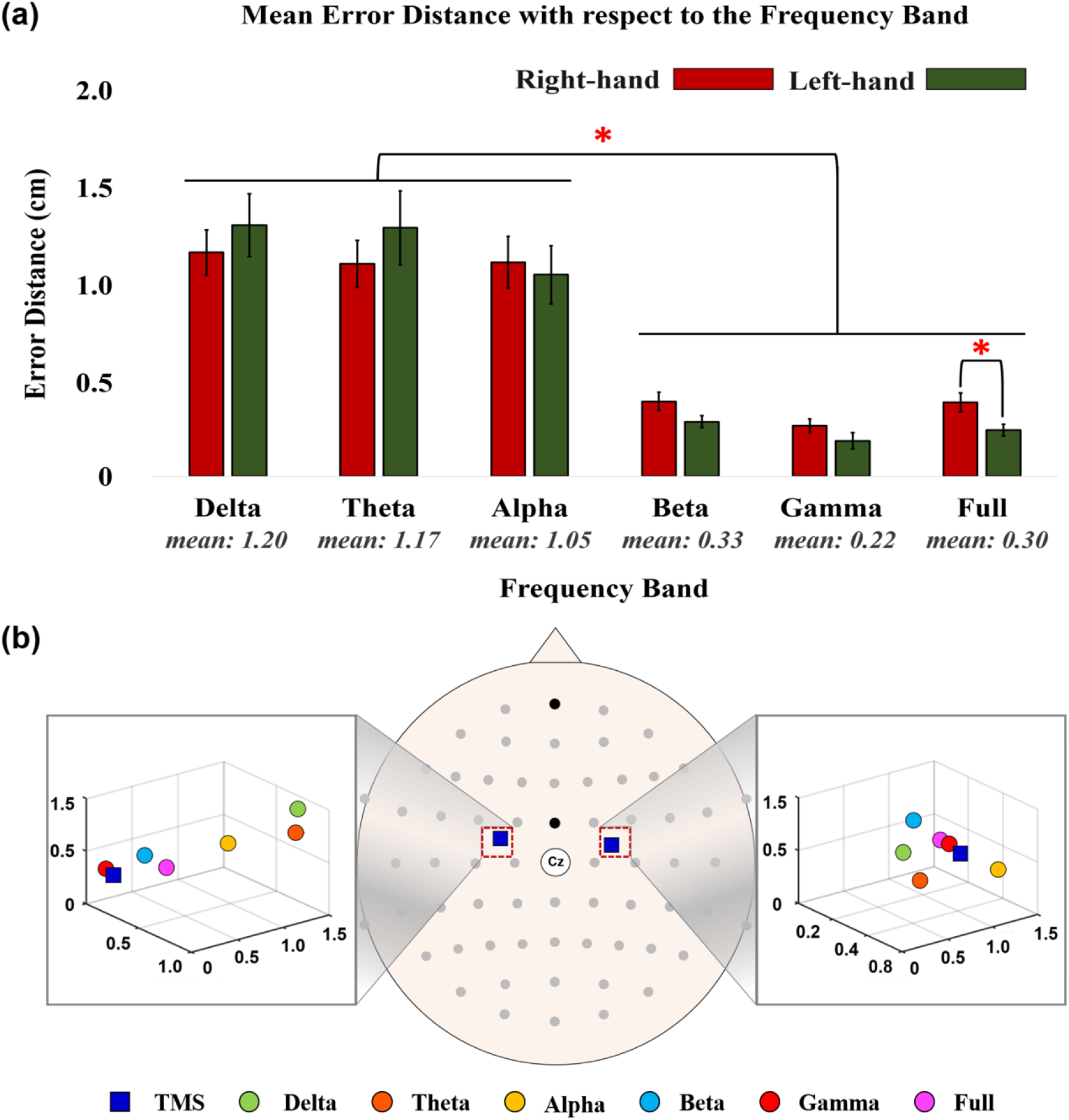
(a) Mean error distances between the motor hotspot locations identified by TMS-induced MEP and our EEG-based machine learning approach with respect to the frequency band (RM-ANOVA with Bonferroni corrected *p*-value < 0.05: delta = theta = alpha > beta = gamma = full for both hands). The numbers below the bar graphs represent the mean error distances of the left and right hands. No significant difference was observed between the left and right hands for all the frequency bands in terms of the error distance (paired t-test *p* > 0.05), except the full band (right hand > left hand). (b) A representative example showing the 3D locations of the motor hotspots identified by TMS-induced MEP (blue rectangle) and our EEG-based approach with respect to the frequency band.

Figure 5(a) presents the mean error distances of the motor hotspot locations estimated by our EEG-based approach for each hand with respect to the number of channels. In general, the mean error distance monotonically and significantly increased as the number of channels decreased (RM-ANOVA with Bonferroni corrected *p*-value < 0.05: Ch_Set5 > Ch_Set4 > Ch_Set3 = Ch_Set2 > Ch_Set1 for the right hand and Ch_Set5 > Ch_Set4 = Ch_Set3 > Ch_Set2 > Ch_Set1 for the left hand). No significant difference was observed between the left and right hand for all channel sets in terms of the error distance (paired t-test *p* > 0.05), except Ch_Set3 (left hand > right hand). Figure 5(b) shows the 3D locations of the motor hotspots identified by TMS and our EEG-based approach with respect to the number of channels, showing a similar trend to the result shown in Figure 5(a).

**Figure 5.**
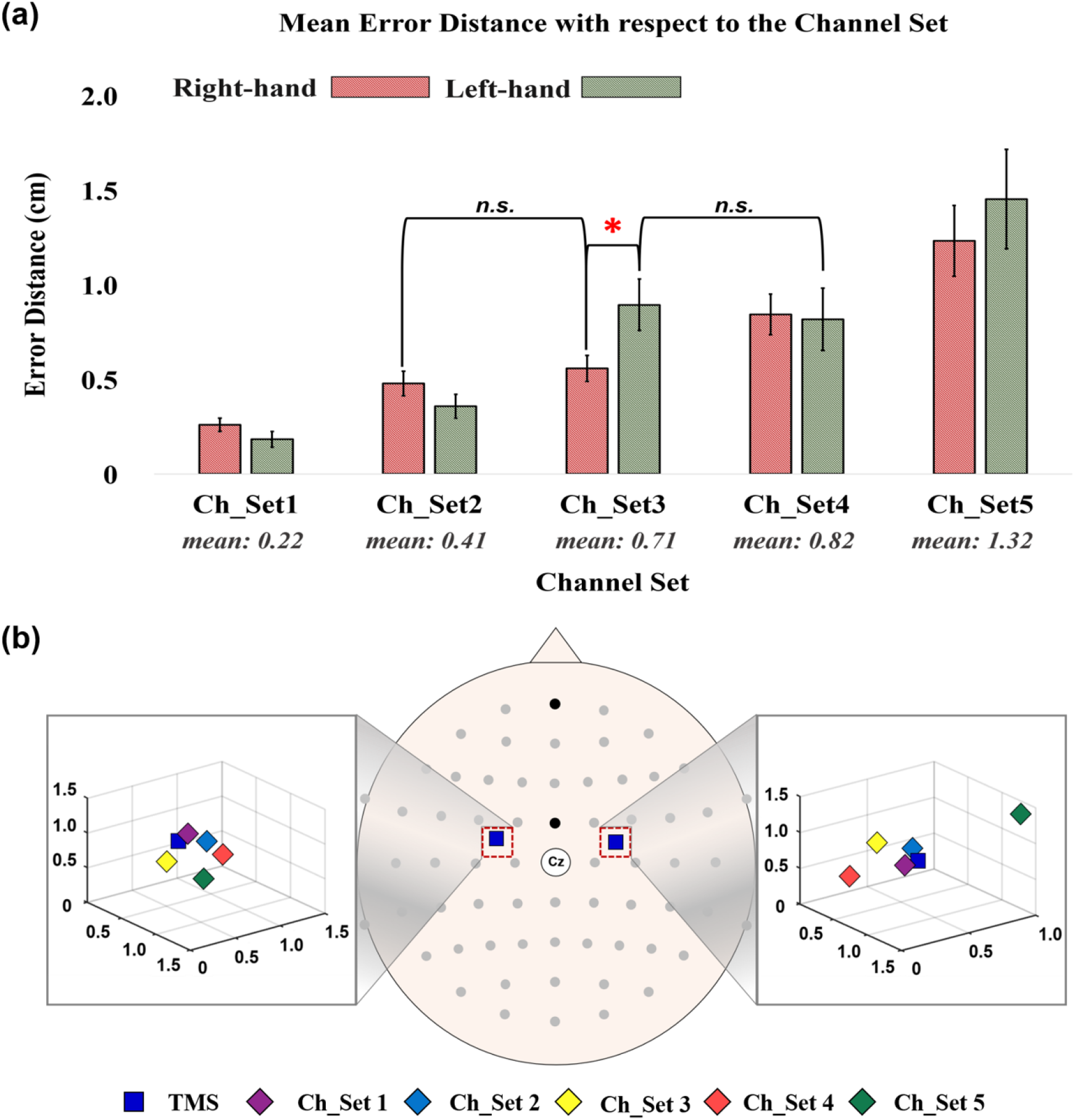
(a) Mean error distances between the motor hotspot locations identified by TMS-induced MEP and our EEG-based machine learning approach with respect to the number of channels (RM-ANOVA with Bonferroni corrected *p*-value < 0.05: Ch_Set5 > Ch_Set4 > Ch_Set3 = Ch_Set2 > Ch_Set1 for the right hand and Ch_Set5 > Ch_Set4 = Ch_Set3 > Ch_Set2 > Ch_Set1 for the left hand). The numbers below the bar graphs represent the mean error distances of the left and right hands. No significant difference was observed between the left and right hand for all channel sets in terms of the error distance (paired t-test *p* > 0.05), except Ch_Set3 (left hand > right hand). The abbreviation, *n*.*s*., means no significant difference. (b) A representative example showing the 3D locations of the motor hotspots identified by TMS (blue rectangle) and the EEG-based approach with respect to the number of channels.

## Discussion

In this study, we proposed a novel EEG-based motor-hotspot-identification method using an ANN as a potential alternative to TMS to determine a tES target location for motor rehabilitation. A minimum mean error distance of 0.22 ± 0.03 cm was attained when using the gamma band PSD information extracted using all 63 channels, demonstrating the proof-of-concept of the proposed EEG-based motor-hotspot-identification method. An EEG device is required for the use of the proposed motor-hotspot-identification approach based on a machine learning technique. In recent years, a portable tES device integrated with an EEG device was introduced (e.g., NeuroElectronics Starstim), which could facilitate the use of our motor-hotspot-identification approach without using TMS.

To check the practical feasibility of the proposed motor-hotspot-identification method, the effect of the number of channels was investigated using the gamma band PSD features. As expected, the lowest mean error was obtained using all channels, and the mean error distance increased as the number of channels decreased (Figure 5). We obtained a mean error distance of less than 1 cm with a number of channels above 17 (Ch_Set4) broadly attached on the motor cortex. In addition, a mean error distance of approximately 1.32 cm was found when using only nine channels attached on the midline of the motor cortex. It was documented that reliable FDI MEP was observed with an area of approximately 12.9 cm^2^ (3.6 cm × 3.6 cm), meaning that a reliable MEP could be evoked with a distance of up to approximately 1.8 cm based on the center of the motor hotspot area [29]. Therefore, although the mean error distance of our proposed motor-hotspot-identification method statistically increased as the number of channels decreased, it is expected that our EEG-based machine-learning approach could be utilized by employing only the motor cortex channels for motor hotspot identification, thereby improving its practicality.

The performance of our proposed motor-hotspot-identification method was better when using higher frequency EEG features (delta: 1.20 ± 0.09 cm; theta: 1.17 ± 0.10 cm, alpha: 1.05 ± 0.09 cm, beta: 0.33 ± 0.03 cm, gamma: 0.22 ± 0.03 cm, full: 0.30 ± 0.03 cm). In particular, beta and gamma PSDs showed significantly lower mean error distances than those of delta, theta, and alpha PSDs. Much evidence has been accumulated indicating that a hand motor task significantly changes EEG frequency information in relatively higher frequency bands, i.e., alpha, beta, and gamma bands [30–34], and thus the higher performance obtained using higher frequency EEG features can be explained from a neurophysiological point of view. However, alpha PSDs did not show better performance as compared to delta and theta PSDs even though alpha band is also closely associated with motor tasks, which should be further investigated in future studies to optimize EEG frequency bands for more accurate motor hotspot identification based on the proposed EEG-based machine-learning approach.

The difference between the mean error distances for the left and right hands were not statistically significant for most comparison cases; two cases showed significant difference (full band and Ch_Set3). This might indicate that our proposed approach is not sensitive to handedness for motor hotspot identification. However, as all subjects recruited in this study were right-handed, additional experiments are required with left-handed subjects to carefully address the mentioned hypothesis.

The application of tES is not only limited to motor rehabilitation but can also be applied to various psychiatric disorders, such as depression, schizophrenia, and attention deficit hyperactivity disorder, to improve not only their cognitive functions, but also neurologically relieve their symptoms [35–39]. In order to apply tES to psychiatric disorders, a target location should be first determined, similarly to motor hotspots for motor rehabilitation. For cognitive rehabilitation, in general, the anodal electrode is attached to F3 according to the international 10-20 system to stimulate the dorsolateral prefrontal cortex (DLPFC), which is known to be associated with various cognitive functions, and the cathodal electrode is attached to F4 or the supraorbital area of the contralateral hemisphere [39–41]. However, as the location of motor hotspots varies from an individual to another, it could be assumed that the tES target location for cognitive rehabilitation is also slightly different between individuals. Therefore, our proposed motor-hotspot-identification approach might be also used for precisely finding a tES target location to maximize the positive effect of cognitive rehabilitation.

On the other hand, the proposed motor-hotspot-identification method used the motor hotspot coordinates identified by TMS as an input to construct the ANN model. Considering the practical use of our proposed method without TMS, the motor hotspot information obtained using TMS should be excluded in the process of finding a motor hotspot using a machine learning technique, and thus we will continuously advance our proposed algorithm in such a way that the motor-hotspot-identification algorithm ultimately does not require the motor hotspot information obtained using TMS for a new subject. Despite the mentioned limitation, it is thought that the results shown in this study could prove the proof-of-concept of the proposed EEG-based motor-hotspot-identification method.

## Conclusion

In this study, we proposed a novel EEG-based motor-hotspot-identification approach as an alternative to TMS and demonstrated its feasibility via EEG experiments. We also confirmed the possibility of using our proposed method to the development of a practical EEG-based motor-hotspot-identification system with a small number of channels attached only on the motor cortex. Because the brain activity patterns of patients with motor impairment would be different from those of healthy subjects, our proposed motor-hotspot-identification method should be further verified with patients to carefully demonstrate its clinical feasibility.

## Abbreviations

tES: transcranial electrical stimulation
NIBS: non-invasive brain stimulation
tDCS: transcranial direct current stimulation
tACS: transcranial alternating current stimulation
tRNS: transcranial random noise stimulation
TMS: transcranial magnetic stimulation
ADHD: attention deficit hyperactivity disorder
DLPFC: dorsolateral prefrontal cortex
MEP: motor evoked potential
EEG: electroencephalography
IRB: institutional review board
FDI: first dorsal interosseous
EMG: electromyography
ICA: independent component analysis
PSD: power spectral density
FFT: fast Fourier transform
ANN: artificial neural network
MRCP: movement-related cortical potential.

## Acknowledgements

Not applicable.

## Authors’ contributions

GYC, NJP, WSK, and HJH participated in designing the experimental paradigm, interpreted the data, and drafted the manuscript. GYC and HTL conducted data collection, and GYC and CHH analyzed the data.

## Availability of data and material

All data measured and analyzed during this study will be published in a public repository after official publication.

## Ethics approval and consent to participate

This study was approved by the Institutional Review Board (IRB) of Kumoh National Institute of Technology (No. 6250) and was conducted in accordance with the principles of the declaration of Helsinki.

## Consent for publication

cNot applicable. No subject is identifiable by the data presented in the figures and tables.

